# Boron Hyperaccumulation Mechanisms in *Puccinellia distans* as Revealed by Transcriptomic Analysis

**DOI:** 10.1101/110403

**Authors:** Saniye Elvan Öztürk, Mehmet Göktay, Canan Has, Mehmet Babaoğlu, Jens Allmer, Sami Doğanlar, Anne Frary

## Abstract

Boron is an essential plant micronutrient; but is toxic at high concentrations. Boron toxicity can severely affect crop productivity in arid and semi-arid environments. *Puccinellia distans (Jacq.) Par1.*, common alkali grass, is found throughout the world and can survive under boron concentrations that are lethal for other plant species. In addition, *P. distans* can accumulate very high levels of this element. Despite these interesting features, very little research has been performed to elucidate the boron tolerance mechanism in this species. In this study, *P. distans* samples were analyzed by RNA sequencing to identify genes related to boron tolerance and hyperaccumulation. Abundance levels of selected differentially expressed transcripts were validated by real-time PCR. The results indicated that the hyperaccumulation mechanism of *P. distans* involves many transcriptomic changes including those that lead to: alterations in the malate pathway, changes in cell wall components that allow sequestration of excess boron without toxic effects, and increased expression of at least one putative boron transporter and two putative aquaporins. Elucidation of the boron accumulation mechanism is important to develop approaches for bioremediation of boron contaminated soils.

## 1. INTRODUCTION

Boron (B) is a plant micronutrient and is essential for plant cell wall formation (Marschner &Marschner, 2012). Boron plays roles in many physiological and biochemical pathways for example in growth and development, including metabolism of nucleic acids, carbohydrates, proteins and indole acetic acid. Cell wall synthesis and structure (Matoh, 2001) as well as membrane integrity and function are further aspects boron is associated with (Goldbach et al., 2001; Goldbach H. E, 1997; Marschner, 2011). Soil boron concentrations vary from deficient to toxic in different regions of the world (Yau et al., 1995). The majority of soil boron is found in Turkey (72.5 % of world reserves), followed by Russia, USA, and China (BOREN, 2012). Important boron reserves are located in the northwest part of Turkey in Balikesir, Bursa, Kutahya and Eskisehir provinces (Ozturk et al., 2010). Boron levels are also affected by rainfall and boron-rich groundwater which can cause boron toxicity in arid and semi-arid regions of the eastern Mediterranean, western Asia/north Africa, the Indian subcontinent, China, Japan and South America (Nable et al., 1997; Tanaka and Fujiwara, 2008; Sang et al., 2015).

The levels of boron that cause deficiency and toxicity vary among plants. However, in general, the difference between boron deficiency and toxicity is observed within a very narrow range. Soil boron concentrations less than 0.5 mg kg^−1^ are associated with deficiency while levels greater than 5.0 mg kg^−1^ cause toxicity (Ryan and Rashid, 2006). The tipping point between deficiency and toxicity is even finer when internal boron concentration is considered. For example, soybean leaf elongation was optimal at 10 to 12 mg kg^−1^ dry weight and impaired at both higher and lower concentrations (Kirk and Loneragan, 1988). Boron deficiency has damaging effects on many biological processes such as carbohydrate metabolism, legume nitrogen fixation (Yamagishi and Yamamoto, 1994), plasma membrane structure and cell wall integrity (Camacho-Cristóbal et al., 2008). Boron toxicity causes decreases in crop growth and productivity (Ozturk et al., 2010; Yau et al., 1995). Boron toxicity also disrupts metabolism by altering cell division and development as well as reducing cell growth. Symptoms of boron toxicity are typically observed on mature leaf margins which become chlorotic and/or necrotic (Ozturk et al., 2010).

Although excess boron is usually toxic, some plant species can tolerate high concentrations. One such species, *Puccinellia distans* (weeping alkali grass) was found to be extremely tolerant to boron toxicity (Padmanabhan et al., 2012). Under hydroponic conditions, *P. distans* shoots accumulated approximately 6000 mg kg^−1^ (Bar, 2015). Recent work by Ramila et al. (2016) suggested that a related species, *Puccinellia frigida,* is also highly tolerant to boron toxicity with approximately 4000 mg kg^−1^ accumulation in shoots (Rámilaa et al., 2015; Rámila et al., 2016). As a result, Puccinellia species are considered to be important candidates for phytoremediation applications. Phytoremediation is a more practical and promising solution for reducing the boron content of arid and semi-arid soils compared to chemical, physical and biological boron-removal technologies which are expensive and impractical (Padmavathiamma and Li, 2007). Despite its potential for phytoremediation, the boron hyperaccumulation mechanism of Puccinellia has not yet been elucidated.

The first boron transport mechanism described in plants was passive transport of uncharged boric acid across the plasma membrane (Marschner and Marschner, 2012). To understand the genetic factors involved in boron transport, early studies investigated nuclear genes in bread wheat and determined that the *Bo1*, *Bo2*, and *Bo3* nuclear genes control boron tolerance (Paull et al., 1992). The *BoT1* and *BoT2* genes were then identified in barley (Jenkin, 1993) and durum wheat (Jamjod, 1996). Later studies indicated that these genes encode active boron transporter proteins. These findings were further supported by the fact that boron uptake was hindered by mutations in BOR transporter genes and metabolic inhibitors in species including *A. thaliana* (Takano et al., 2001), barley (Hayes, 2004), sunflower (Matoh and Ochiai, 2005), wheat (Reid, 2007), and rice (Uraguchi and Fujiwara, 2011). Boron uptake is also mediated by major intrinsic channel-like transporter proteins (MIPs) in sunflower (Dannel et al., 2000), squash (Dordas & Brown, 2000), and Arabidopsis (Takano et al., 2006). Among these MIPs, a number of aquaporins are reported to be permeable to physiologically important molecules such as boron, silicon, ammonia, hydrogen peroxide, and carbon dioxide (Chaumont et al., 2005). Indeed, two aquaporins in *A. thaliana* were shown to have boric acid channel activity (Takano et al., 2006;Takano et al., 2008).

The main goal of this study was to examine the hyperaccumulation/tolerance mechanisms of *P. distans* under high boron conditions. To achieve this goal, we used RNA sequencing to compare the expression profiles of plants grown under control and normally toxic boron conditions. Differentially expressed transcripts were annotated at the protein level and gene ontology and pathway analyses were performed against the model organisms, *O. sativa* and *A. thaliana*. The differentially regulated proteins were found to be located in several areas including membrane-associated complexes, the cytosol and nucleus. These proteins play roles in several molecular pathways including carbon, energy, amino acid, and lipid metabolism; signal transduction and transport. In this way, we identified candidates including an *A. thaliana* boron transporter, an *O. sativa* nodulin-26 like Intrinsic Protein (NIP) and other stress response-related transcripts in *P. distans*. A greater understanding of the hyperaccumulation/tolerance mechanism(s) of the monocot *P. distans* will provide insight into ways of coping with boron stress toxicity in agriculturally important cereals such as rice, maize, and sorghum. Thus, while *P. distans* itself might be used to eliminate excess boron from soil, the tolerance mechanism(s) could be transferred to crop plants to confer boron tolerance and/or to develop a faster-growing, higher biomass phytoremediation system.

## 2. MATERIALS & METHOD

### 2.1. Plant material and boron treatment

Seeds of *P. distans* collected from a boron mining site, Kirka-Eskisehir, Turkey (39° 17′ 23.7156” and 30° 31′ 33.4812”) (Babao□lu et al., 2004), were germinated in potting soil. The germinated seedlings were grown for 2 weeks in a growth chamber maintained at 25 ± 2°C, 60 % relative humidity and 16 h photoperiod (Stiles et al., 2010). After four weeks, plants were transferred to half strength Hoagland solution (Hoagland and Arnon, 1950) and grown for one week. Plants were separated into two groups with three replicates: 0.5 mg L^−1^ boron was applied to the control group and 500 mg L^−1^ was applied to the stress group (Stiles et al., 2010). The boron concentration in Hoagland solution was adjusted with boric acid and the solution was changed once every three days for three weeks. After 3 weeks of treatment, plants were removed from the Hoagland solution and rinsed of any contamination with RNase-free water. The shoot tissues were then frozen in liquid nitrogen and stored at −80°C.

### 2.2. RNA isolation

Total RNA was isolated from the shoot samples using an RNeasy Plant Mini Kit (Qiagen, Maryland, USA). The quality and quantity of isolated shoot RNA from control and stress samples were measured using a Nanodrop ND-100 device (Nanodrop Technologies, Wilmington, DE, USA).

### 2.3. RNA sequencing and analysis

Extracted total RNA samples were processed using a TruSeqTM RNA Sample Preparation Kit (Illumina, Tokyo, Japan) for cDNA library construction and subsequent EST identification. ESTs were sequenced for control and stress libraries using an Illumina High-Seq 2000 platform (Takara, Tokyo, Japan) to generate 125 bp paired end (PE) reads. RNA sequencing was performed by GATCBiotech (Constance, Germany). Raw data consisted of 45 million reads for each sample with read length fixed at 125 nucleotides.

The Cutadapt2 (version 1.9.1) program (Martin, 2011) was employed with default parameters to remove adapter sequences and low quality nucleotides from raw reads. Reference transcriptome construction was performed using the Trinity (version 2.2.0) assembly tool (PMID: 23845962) (Haas et al., 2013). All reads from the control and stress samples were treated the same. The reference transcriptome was used to map the cleaned reads from control and stress datasets individually using the Bowtie2 program (version 2.1.0) (Langmead & Salzberg, 2012). Mapping information for each dataset was saved in a binary alignment information (bam) file. The bam files were used as inputs for differential gene expression analysis performed with the Cufflinks (version 2.2.1) pipeline (Cufflinks, Cuffmerge, and Cuffdiff) (Trapnell et al., 2012). Results were analyzed and visualized using the cummeRbund (version 2.15) R statistics package (Goff et al., 2013). Differentially expressed gene sequences were extracted based on q-value (q-value threshold < 0.05, where q-value is the false discovery rate adjusted p-value of the test statistic). Differentially expressed candidates were then annotated using Blast2GO (version 4.0.7) (Conesa et al., 2005) against a custom protein database which included *O. sativa* and *A. thaliana* proteins in UniProt Knowledgebase (release date: November 2016). Gene ontology analysis was carried out with QuickGO (Binns et al., 2009). Protein functional classification based on gene ontology was performed by PANTHER Protein Classification System (Accession date: November, 2016) (Mi et al., 2016). Annotated transcripts were search against the KEGG (Kyoto Encyclopedia of Genes and Genomes) database (Tanabe & Kanehisa, 2012) to reveal the pathways in which up and down-regulated transcripts have roles.

### 2.4. Evolutionary conservation

Phylogenetic analyses of boron transporters and other ion transporters of multiple species were performed to determine their sequence similarity and evolutionary conservation. This analysis included all of the fourteen transporters found to be up or down-regulated by boron stress in this work: *O. sativa* boron NIP transporter (UniProtID: Q949A7), *A. thaliana* TIP transporter (UniProtID: O82598), *A. thaliana* BOR6 transporter (UniProtID: Q3E954), *A. thaliana* sugar transporters (UniProtIDs: Q4F7G0, O04249), *O. sativa* sulfate transporter (UniProtID: Q8S317), *O. sativa* anion transporter (UniProtID: Q652N5), *A. thaliana* inorganic phosphate transporter (UniProtID: Q38954), and *A. thaliana* ABC transporters (UniProtIDs: Q9C9W0, Q8LPJ4, Q7GB25, Q9FWX7, Q9FJH6, Q9M1H3). In addition, 11 boron transporters described in the literature were used in the analysis including those from *O. sativa* (UniProtIDs: Q1ZYR7, Q7X9F3), *A. thaliana* (UniprotID: Q8VYR7, Q9XI23, Q93Z13, Q9M1P7, Q9SUU1, Q9SSG5), *Hordeum vulgare* (UniProtID: M0Z9M9), *Brassica napus* (UniProtID: D5LGA1, D5LG97). Two *A. thaliana* NIP transporters (UniProtID: Q9SV84, Q9SAI4) were also included. Sequences for these proteins were retrieved from the UniProtKB database. *A. thaliana* nuclear transport factor 2 (NTF2) protein (UniprotID: F4J8X6) was included as outgroup. Multiple sequence alignment was conducted with the default settings of ClustalW (Larkin et al., 2007). All positions with continuous alignment gaps were eliminated using the MEGA 7.0 suite (Tamura et al., 2013). Phylogenetic tree construction was performed using the maximum likelihood method based on the Le Gascuel substitution model (Le & Gascuel, 2008). The Gamma distribution was used to model evolutionary rate differences among sites (with default parameters except for G which was set to 13, 3340). Bootstrap values were inferred from 1000 replicates. In order to compute phylogenetic distances, the boron and other transporters were separated into two groups. Within and between groups distances were calculated by computing the number of amino acid differences per site, averaged over all sequence pairs.

### 2.5. Quantitative real-time PCR

The expression of 30 genes that responded to boron stress in *P. distans* shoots was confirmed by qRT-PCR (quantitative real-time polymerase chain reactin) analysis using three technical replicates from one of the three biological replicates used for RNA-seq analysis. The extracted total RNA was treated with DNase I (Takara, Shiga, Japan). GoTaq^®^ 2-Step RT-qPCR kit (PROMEGA) was used for cDNA synthesis and the resulting cDNAs were amplified in the LightCycler^®^ 480 system (Roche, Basel, Switzerland) using transcript-specific primers (Suppl. Table 1). The 30 transcripts were selected randomly from the top 50 up and down-regulated transcripts. In addition, primers were used for one known ABC transporter and one known boron transporter (Padmanabhan et al., 2012).

**Table 1.** Summary of transcripts/genes statistics of *P. distans*.

| Parameter |  |
| --- | --- |
| Number of Reads (Control + Stress) | 179,792,482 |
| Average Raw Read Length (Control + Stress) | 125 nt |
| Average Contig Length | 496.73 nt |
| Median Contig Length | 317 nt |
| N50 | 609 nt |
| Total Assembly Size | 284,371,845 nt |

## 3. RESULTS

### 3.1. Analysis of the *P. distans* transcriptome with RNA-seq

To investigate the transcriptomic response of *P. distans* to boron stress, four cDNA libraries were generated from two replicate mRNA samples from boron-treated and control shoot samples. These libraries were sequenced using Illumina deep-sequencing HiSeq™ 2000. Raw reads of control (90,514,028) and boron-treated stress (89,278,454) samples were generated. Raw reads were cleaned of adapter sequences using the Cutadapt2 tool resulting in 88,663,592 (98.0 %) and 82,494,636 (92.4 %) reads for control and stress samples, respectively.

The cleaned reads from the two samples were treated as input for the Trinity assembler. All reads were mapped to the constructed reference transcriptome individually using the Bowtie2 program. Mapping ratios for control and stress samples were 97.8 % and 97.3 % of cleaned reads, respectively.

Based on transcripts that mapped to the reference transcriptome, 284,371,845 bases were assembled from 179,792,482 reads (control + stress) (Table 1). Median and average contig lengths were 317 and ~497 nt, respectively (Table 1). Additionally, the N50 value indicated that at least half the assembled bases were found in contigs that were at least 609 nt long (Table 1).

### 3.2. Differential gene expression between control and stress conditions

Differentially expressed transcripts were detected using the RNA-Seq alignments and Cufflinks processing. Cuffdiff tracked the mapped reads and determined the fragments per kb per million mapped reads (FPKM) for each transcript in all samples. These FPKM values were used to detect up and down-regulated transcripts and most differential transcripts were found to be present or absent under control or stress conditions (Figure 1). A total of 2242 of 3312 differentially expressed transcripts were up-regulated (67.7 %) while 1070 of 3312 (32.3 %) were down-regulated (Suppl. Table 2).

**Figure 1.**
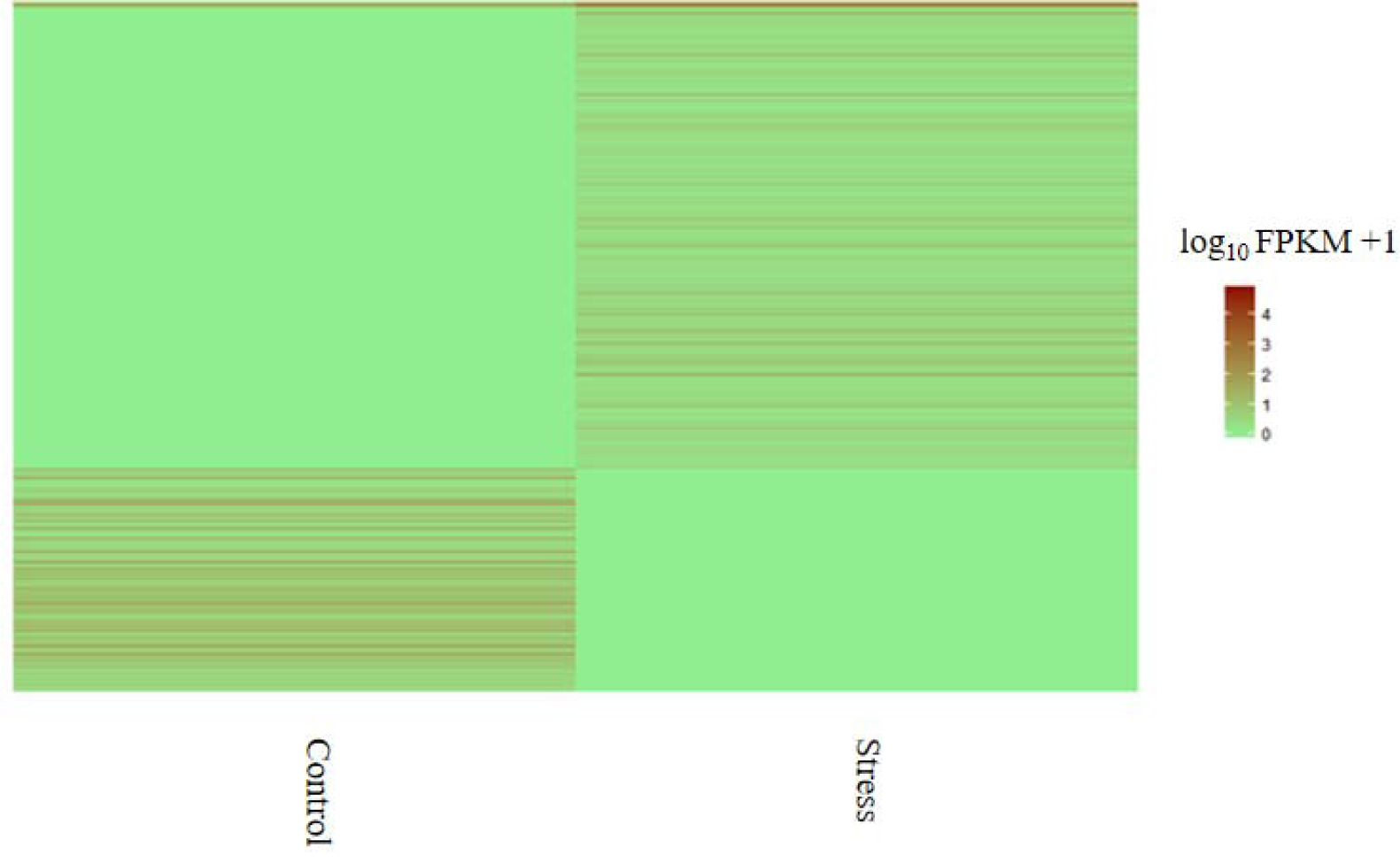
Heat map of log transformed FPKM values of 3312 differentially expressed transcripts from RNA-Seq of control and stress samples. FPKM ranges from 0 (green) to 5 (red).

**Table 2:** Transcripts with high differential expression based on FPKM value. Annotated UniProt ID, sequence description and organism [*A. thaliana* (At) and *O. sativa* (Os)] are included.

| UniProtID/ Annotation | Organism | Transcript Name | Difference* |
| --- | --- | --- | --- |
| F4I600_ARATH/ V-ATPase-related | At | TRINITY_DN213899_c1_g5_i1:0-169 | 20561.12 |
| H32_ARATH/ Histone | At | TRINITY_DN99923_c0_g1_i1:0-171 | 845.90 |
| RRAA2_ARATH/ 4-hydroxy-4-methyl-2-oxoglutarate aldolase 2 | At | TRINITY_DN220021_c1_g4_i1:0-584 | 426.28 |
| PSBR_ARATH/Photosystem II 10 kDa chloroplastic | At | TRINITY_DN192927_c1_g1_i1:132-532 | 329.61 |
| RD22_ARATH/ BURP domain RD22 | At | TRINITY_DN131860_c0_g2_i1:0-222 | 315.95 |
| RRAA3_ARATH/ 4-hydroxy-4-methyl-2-oxoglutarate aldolase 3 | At | TRINITY_DN220021_c1_g4_i2:0-391 | 299.33 |
| RRAA3_ARATH/ 4-hydroxy-4-methyl-2-oxoglutarate aldolase 3 | At | TRINITY_DN214561_c1_g18_i1:0-365 | 253.61 |
| H2B10_ARATH/ Histone | At | TRINITY_DN199272_c0_g3_i1:2-246 | 168.98 |
| H2B1_ARATH/ Histone | At | TRINITY_DN150574_c0_g1_i1:0-201 | 130.08 |
| A0A160DR54_ORYSA/ Dirigent (Fragment) | Os | TRINITY_DN215985_c1_g3_i1:4-592 | 126.38 |
| RL40B_ARATH/ Ubiquitin-60S ribosomal L40-2 | At | TRINITY_DN67858_c0_g1_i1:0-219 | 85.64 |
| H2A1_ARATH/ Probable histone | At | TRINITY_DN28983_c0_g1_i1:38-250 | 80.83 |
| FRO7_ARATH/ Ferric reduction oxidase chloroplastic | At | TRINITY_DN224067_c1_g1_i1:0-1399 | 79.87 |
| H32_ARATH/ Histone | At | TRINITY_DN271392_c0_g1_i1:0-479 | 66.13 |
| H2B3_ARATH/ Histone | At | TRINITY_DN199272_c0_g1_i1:0-245 | 65.63 |
| H2A1_ARATH/ Probable histone | At | TRINITY_DN113560_c0_g1_i1:0-238 | 63.07 |
| XTH22_ARATH/ Xyloglucan endotransglucosylase hydrolase 22 | At | TRINITY_DN221642_c3_g12_i1:2-232 | 57.72 |
| H32_ARATH/ Histone | At | TRINITY_DN48466_c0_g1_i1:30-384 | 51.59 |
| Q01KB9_ORYSA/ Autophagy-related | Os | TRINITY_DN202627_c0_g7_i1:92-353 | 45.84 |
| ACT3_ARATH/ Actin-3 | At | TRINITY_DN272586_c0_g1_i1:1-261 | 39.90 |
| SYHM_ARATH/ Histidine-tRNA chloroplastic mitochondrial | At | TRINITY_DN179460_c0_g1_i1:0-227 | 39.49 |
| COX3_ARATH/ Cytochrome c oxidase subunit 3 | At | TRINITY_DN264216_c0_g1_i1:1-257 | 38.02 |
| H4_ARATH/ Histone H4 | At | TRINITY_DN8209_c0_g1_i1:0-419 | 31.53 |
| Q949A7_ORYSA/ aquaporin (Fragment) | Os | TRINITY_DN210823_c0_g3_i1:4-269 | 30.12 |
| H2A1_ARATH/ Probable histone | At | TRINITY_DN202443_c4_g5_i3:0-460 | 29.44 |
| FRS5_ARATH/ FAR1-RELATED SEQUENCE 5 | At | TRINITY_DN114402_c0_g1_i1:9-274 | 29.40 |
| H2A1_ARATH/ Probable histone | At | TRINITY_DN202443_c4_g4_i2:0-314 | 25.88 |
| H2A1_ARATH/ Probable histone | At | TRINITY_DN202443_c4_g4_i3:0-340 | 25.53 |
| Q9LK47_ARATH/ AT3g23700 MYM9_3 | At | TRINITY_DN219995_c1_g16_i1:0-256 | 25.08 |
| A0A0K0WRF9_ORYSA/ NBS-LRR type R | Os | TRINITY_DN226626_c0_g8_i1:3-253 | 24.76 |
| H2A1_ARATH/ Probable histone | At | TRINITY_DN202443_c4_g1_i1:0-255 | 24.25 |
| NEN3_ARATH/ NEN3 | At | TRINITY_DN226592_c4_g15_i1:8-257 | 21.50 |
| RL341_ARATH/ 60S ribosomal L34-1 | At | TRINITY_DN295721_c0_g1_i1:0-414 | 20.41 |
| F4HS76_ARATH/ Glycerol kinase | At | TRINITY_DN140573_c0_g2_i1:0-250 | 20.29 |
| RL142_ARATH/ 60S ribosomal L14-2 | At | TRINITY_DN179003_c1_g2_i1:7-283 | 20.12 |
| RL74_ARATH/ 60S ribosomal L7-4 | At | TRINITY_DN294520_c0_g1_i1:0-828 | 20.02 |
| F4IJE1_ARATH/ Phox domain-containing | At | TRINITY_DN278982_c0_g1_i1:1-268 | 19.68 |
| COX2_ARATH/ Cytochrome c oxidase subunit 2 | At | TRINITY_DN284184_c0_g1_i1:0-703 | 19.48 |
| Y1457_ARATH/ Acyltransferase chloroplastic | At | TRINITY_DN222292_c3_g2_i1:0-281 | 18.69 |
| Q9LTW8_ARATH/ CTP synthase | At | TRINITY_DN1712_c0_g1_i1:0-285 | 18.65 |
| A0A0U2JFK9_ORYSA/ NBS-LRR-like resistance | Os | TRINITY_DN213110_c2_g2_i4:22-323 | 16.98 |
| UBQ3_ARATH/ Polyubiquitin 3 | At | TRINITY_DN186267_c2_g1_i1:18-638 | 16.88 |
| ACT11_ARATH/ Actin-11 | At | TRINITY_DN204305_c3_g6_i2:0-375 | 16.85 |
| A8MRY7_ARATH/ WD-40 PCN | At | TRINITY_DN224769_c1_g7_i1:4-287 | 15.55 |
| Q9STF8_ARATH/ PR-6ase inhibitor family | At | TRINITY_DN127914_c0_g2_i1:0-296 | 15.37 |

\*Difference: FPKM (Fragments per Kilobase of Exon per Million Fragments Mapped) difference between control and stress conditions.

### 3.3. Annotation of differentially expressed transcripts

The number of *P. distans* proteins and annotations available in public data repositories is limited. Therefore, *P*. *distans* annotation was conducted using the protein databases of two model organisms: *A. thaliana* and *O. sativa.* Both of these have extensive protein sequence resources available in the UniProt knowledgebase repository. In addition, *O. sativa* and *P. distans* belong to the same genus, Gramineae, indicating a close genetic relationship (Wang et al., 2007). 3312 transcripts were found to be differentially expressed and of these 1652 (49.9%) mapped to 1107 proteins. Multiple transcripts (a total of 335 transcripts) matching 36 proteins were labelled as conflicting due to matching both up and down-regulated transcripts according to their FPKM values. In order to resolve this, BLAST results were inspected manually. In this way, 36 transcripts with the highest sequence identity and best hit score were kept and the remaining transcripts were discarded as non-annotated. Among the annotated transcripts, 28 matched 21 transporter proteins (Suppl. Table 3). The annotations for the most differentially expressed transcripts (based on stress and control FPKM values) are listed in Table 2. Transcripts that did not return any match against rice and *A. thaliana* proteins were annotated against the green plant database (Viridiplantae). However, no additional candidate proteins were found at a significant level of match based on alignment length and E-value (data not shown).

**Table 3.** Molecular function main and subcategories which were most related to boron hyperaccumulation in P. distans.

| Molecular Function GOTerm | Percentage of proteins |
| --- | --- |
| **Binding** | 39.00 % |
| ATP binding | 9.90 % |
| Metal ion binding | 4.60 % |
| RNA binding | 3.80 % |
| DNA binding | 3.70 % |
| Zinc ion binding | 3.20 % |
| GTP binding | 1.60 % |
| Translation initiation factor activity | 1.60 % |
| Nucleic acid binding | 1.50 % |
| Copper ion binding | 1.40 % |
| Transcription factor activity, sequence-specific DNA binding | 1.10 % |
| ADP binding | 1.00 % |
| Heme binding | 1.00 % |
| Calcium ion binding | 0.60 % |
| Iron ion binding | 0.50 % |
| Magnesium ion binding | 0.30 % |
| HSP90 protein binding | 0.20 % |
| 4 iron, 4 sulfur cluster binding | 0.20 % |
| Cobalt ion binding | 0.10 % |
| Manganese ion binding | 0.10 % |
| 2 iron, 2 sulfur cluster binding | 0.10 % |
| Four-way junction DNA binding | 0.10 % |
| Calmodulin binding | 0.10 % |
| Quinone binding | 0.10 % |
| Iron-sulfur cluster binding | 0.10 % |
| **Catalytic activity** | **67.00 %** |
| Structural constituent of ribosome | 4.90 % |
| Protein serine/threonine kinase activity | 1.30 % |
| Protein kinase activity | 1.00 % |
| GTPase activity | 0.90 % |
| ATP-dependent RNA helicase activity | 0.80 % |
| ATPase activity | 0.80 % |
| NADH dehydrogenase (ubiquinone) activity | 0.60 % |
| Proton-transporting ATP synthase activity, rotational mechanism | 0.60 % |
| Structural molecule activity | 0.60 % |
| Oxidoreductase activity | 0.60 % |
| Kinase activity | 0.50 % |
| Signal transducer activity | 0.30 % |
| Peroxidase activity | 0.30 % |
| Metalloendopeptidase activity | 0.30 % |
| Cytochrome-c oxidase activity | 0.30 % |
| Fumarate hydratase activity | 0.20 % |
| Metalloaminopeptidase activity | 0.10 % |
| Calcium-dependent cysteine-type endopeptidase activity | 0.10 % |
| Glutathione transferase activity | 0.10 % |
| Peroxiredoxin activity | 0.10 % |
| Oxidoreductase activity, acting on the CH-CH group of donors, with a flavin as acceptor | 0.10 % |
| Glutathione peroxidase activity | 0.10 % |
| Calcium-transporting ATPase activity | 0.10 % |
| Electron transporter, transferring electrons from COQH2-cytochrome c reductase complex and cytochrome c oxidase complex activity | 0.10 % |
| L-lactate dehydrogenase activity | 0.10 % |
| L-malate dehydrogenase activity | 0.10 % |
| Metallopeptidase activity | 0.10 % |
| Ferric-chelate reductase activity | 0.10 % |
| Substrate-specific transmembrane transporter activity | 0.10 % |
| Oxaloacetate decarboxylase activity | 0.10 % |
| Manganese-transporting ATPase activity | 0.10 % |
| Oxidoreductase activity, acting on paired donors, with incorporation or reduction of molecular oxygen | 0.10 % |
| Malate dehydrogenase (decarboxylating) (NADP+) activity | 0.10 % |
| Oxidoreductase activity, acting on the CH-CH group of donors | 0.10 % |
| Oxalate decarboxylase activity | 0.10 % |
| Oxidoreductase activity, acting on paired donors, with oxidation of a pair of donors resulting in the reduction of molecular oxygen to two molecules of water | 0.10 % |
| L-ascorbate peroxidase activity | 0.10 % |
| Cytochrome-c peroxidase activity | 0.10 % |
| Glutamate decarboxylase activity | 0.10 % |
| Myo-inositol | 0.10 % |
| Oxidoreductase activity, acting on the CH-CH group of donors, NAD or NADP as acceptor | 0.10 % |
| Nuclear export signal receptor activity | 0.10 % |
| Oxidoreductase activity, acting on NAD(P)H, heme protein as acceptor | 0.10 % |
| Aspartate-tRNA ligase activity | 0.10 % |
| Malate dehydrogenase (decarboxylating) (NAD<sup>+</sup>) activity | 0.10 % |
| Oxidoreductase activity, oxidizing metal ions | 0.10 % |
| Protein tyrosine/serine/threonine phosphatase activity | 0.10 % |
| **Transporter activity** | **9.00 %** |
| Transporter activity | 0.40 % |
| Glucose transmembrane transporter activity | 0.20 % |
| Sugar:proton symporter activity | 0.20 % |
| Inorganic phosphate transmembrane transporter activity | 0.20 % |
| Malate transmembrane transporter activity | 0.10 % |
| L-ascorbic acid transporter activity | 0.10 % |
| Secondary active sulfate transmembrane transporter activity | 0.10 % |
| Antiporter activity | 0.10 % |
| Voltage-gated ion channel activity | 0.10 % |
| Anion:anion antiporter activity | 0.10 % |
| Auxin efflux transmembrane transporter activity | 0.10 % |
| Outward rectifier potassium channel activity | 0.10 % |
| Ion transmembrane transporter activity | 0.10 % |
| Ion channel activity | 0.10 % |
| Channel regulator activity | 0.10 % |
| Inorganic anion exchanger activity | 0.10 % |
| Anion transmembrane transporter activity | 0.10 % |
| Inward rectifier potassium channel activity | 0.10 % |
| L-glutamate transmembrane transporter activity | 0.10 % |
| Zinc ion transmembrane transporter activity | 0.10 % |
| Porin activity | 0.10 % |

Functional annotation was performed using Blast2GO and QuickGO based on gene ontology (GO). Cellular component, molecular function, and biological process were analyzed as GO terms for the 1107 annotated proteins. According to the ontology analysis, 906 proteins (81.8 %) were annotated to cellular components. 885 (80.0 %) and 848 proteins (76.6 %) were annotated with their molecular functions and biological processes, respectively.

#### Cellular components

Cellular component analysis lead to the annotation of 906 proteins (81.8 %) to at least one cellular component-associated term. Excess boron is trafficked from the plasma membrane to other organelles such as the vacuole and cell wall to avoid boron toxicity (Hanaoka, Uraguchi, Takano, Tanaka, & Fujiwara, 2014) Thus, only cellular component-associated GO terms that are related to boron hyperaccumulation are presented (Suppl. Table 4). A total of 14.8 % of proteins were associated with the integral component of the membrane, plasma membrane, vacuolar membrane, mitochondrial inner membrane and endoplasmic reticulum membrane. In addition, 1.4 % of proteins were related to the cell wall, 0.8 % were localized in the apoplast and 1.5 % were in the vacuole.

#### Biological processes

Biological process annotations fell into seven main categories: response to stimulus, cellular process, metabolic process, localization, reproductive process, developmental process and signaling (Suppl. Table 5). Proteins related to the response to plant hormones and stress conditions such as salt, various ions, cold, heat, oxidative stress, and water deprivation were identified. Our findings indicated that not only translation and transcription but also protein folding and turnover; cell wall organization and biogenesis; and ion dependent redox homeostasis were regulated under excess boron. Both anabolic processes such as the malate-fumarate pathway and catabolic processes such as protein and lignin pathways were altered under stress conditions. Proteins that play roles in signal transduction, in particular plant hormone-mediated and sugar-mediated signaling pathways were also differentially regulated.

#### Molecular functions

Molecular function annotation resulted in three main categories which contained the most transcripts: transporter activity, catalytic activity and binding (Table 3).

The most abundant molecular function categories were binding activity (39 %) and catalytic activity (67 %). Binding activity included copper, zinc, calcium, manganese, cobalt and ferritin binding indicating that excess boron alters signal transduction and vesicle trafficking in the cell. Catalytic activities of enzymes such as serine/threonine kinase, GTPase, ATPase, H^+^ transporting ATP synthase, and oxidoreductase were regulated under boron stress. Differentially expressed transporters included proton, glucose, malate, inorganic phosphate, L-ascorbic acid, L-glutamate, auxin, and ion transporters. Transporters for minerals including calcium, sulfate, and manganese were also identified in this category. Moreover, three previously identified boron transporters were found: *A. thaliana* BOR6 putative boron transporter protein (Transcript Name: TRINITY_DN188130_c0_g1_i1:0- 1084), *A. thaliana* TIP1-3 protein (Transcript Name: TRINITY_DN222891_c3_g1_i1:0-216), and *O. sativa* NIP protein (Transcript Name: TRINITY_DN210823_c0_g3_i1:4-269).

Annotated transcripts were uploaded to the KEGG system for metabolic pathway analysis (Kanehisa, 2002). A total of 756 proteins were found to have interactions with pathways in the KEGG database (Suppl. Table 6). In this database, 327 (43.25 %) proteins fell into the general metabolism category. Of these metabolism-related proteins, 66 were involved in carbon metabolism, 64 in energy metabolism, 50 in amino acid metabolism, 31 in nucleic acid metabolism, 24 in lipid metabolism and 7 in secondary metabolism. The remaining 34 proteins were placed in other metabolism-related subcategories. A total of 387 proteins (51.2 %) were in the genetic information process category. Of these, the most proteins (184) fell into the translation subcategory. The protein folding category was the second most populous group in genetic information processes with 98 proteins. Environmental information processes were also observed for 152 proteins (20.1 %) with 148 proteins matching signal transduction and four matching membrane transport. Finally, 152 proteins (20.1 %) were associated with cellular processes. In this category, two subgroups resulted in high protein matches: 86 proteins in the cellular growth and death subcategory and 42 proteins in the transport and catabolism subcategory.

### 3.5. Evolutionary relationships among transporters

*O. sativa* and *A. thaliana* are not boron hyperaccumulating plants; therefore, boron transporters in *P. distans* may have been annotated to other ion transporters in these species. Phylogenetic analysis was used to determine the degree of evolutionary similarity between known boron transporters (from *O. sativa, A. thaliana, B. napus,* and *H. vulgare*) and the annotated *A. thaliana* and *O. sativa* boron (BOR6), aquaporin (NIP and TIP), sugar, sulfate, anion, inorganic phosphate, and ABC transporters from our dataset (Figure 2). A total of 28 amino acid sequences were compared over 160 positions. The samples included an *A. thaliana* nuclear transport factor protein as outgroup. The phylogenetic analysis indicated that the identified sugar, sulfate, anion, inorganic phosphate and ABC transporter proteins were distinct from boron transporters which were also distinct from aquaporins. The mean p-distance was 30 % between boron transporters and 60 % between aquaporins.

**Figure 2.**
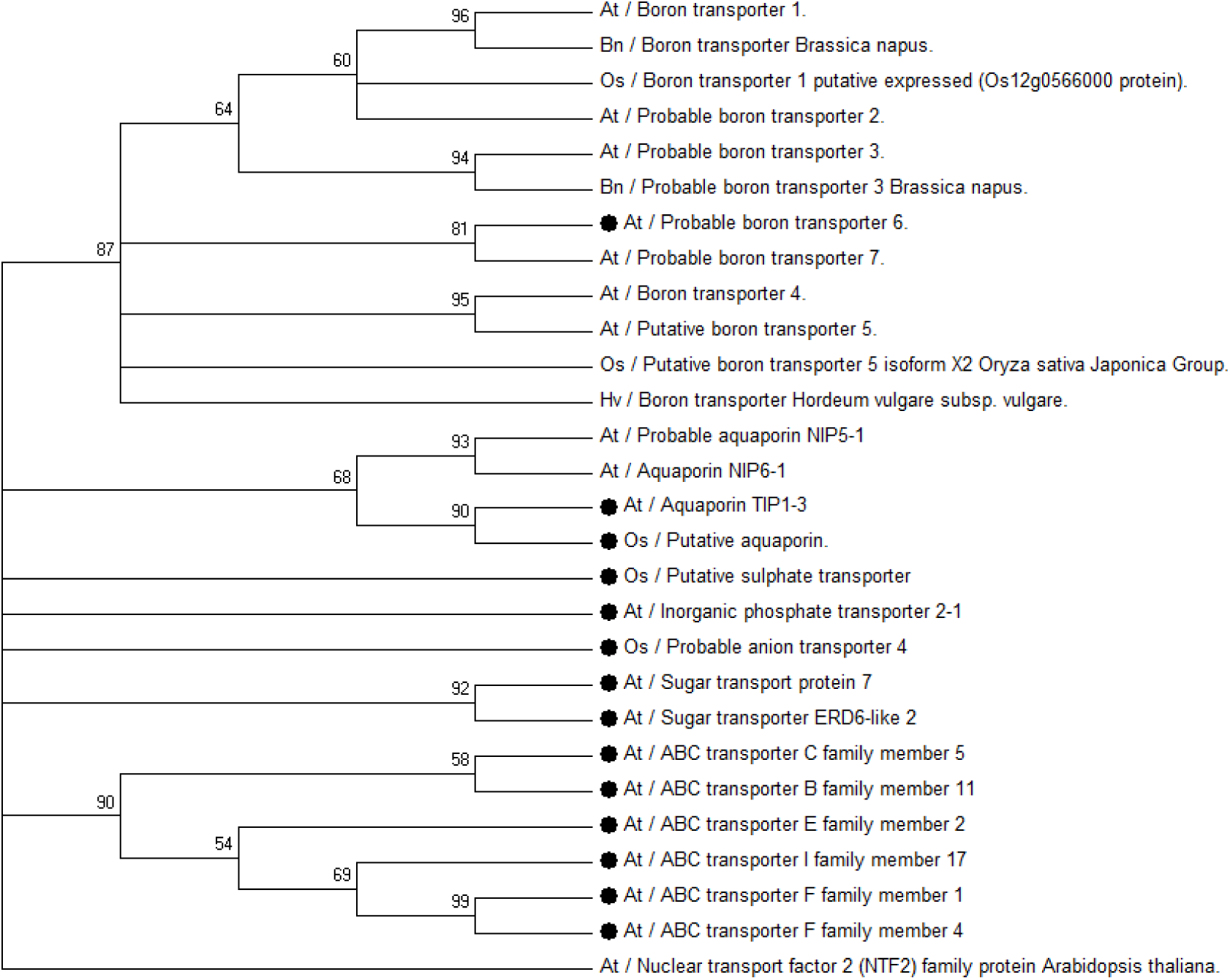
Maximum Likelihood phylogenetic tree representing the evolutionary relationship of boron TIP and NIP transporters and other ion and molecule transporters of several species. *A. thaliana* NTF2 is used as outgroup. The evolutionary history was inferred using the maximum likelihood method based on the Le Gascuel 2008 model. The tree with the highest log likelihood (-8734, 1061) is shown. Bootstrap values are indicated at the branches. There were a total of 160 positions in the final dataset. • indicates the annotated transcripts from this study

#### 3.6. Real-time PCR results of annotated transcripts

A total of 32 transcripts were selected for verification of their expression levels using quantitative-PCR (q-PCR). Transcripts were chosen randomly based on their differential expression between control and stress samples. The q-PCR analysis was performed according to the 2^−ΔΔCt^ method. Up and down-regulated transcripts are shown in Figure 3. The most significant five up-regulated transcripts were annotated to: an aquaporin of *O. sativa* (DN210823); a photosystem II protein in *Arabidopsis* (DN192927); a 4-hydroxy-4-methyl-2- oxoglutarate aldolase in *A. thaliana* (DN220021); a ferric reduction oxidase in *A. thaliana* (DN224067); and FRIGIDA, which is required for FLC (Flowering Locus C) activity in *A. thaliana* (DN111068)(Figure 3A). Significant down-regulation was observed for four transcripts: three of which were annotated to have oxidoreductase activities in *O. sativa* (DN212455, DN205322, DN216505) and one that was annotated to aquaporin TIP1-3 in *A. thaliana* (DN222891)(Figure 3B and Suppl. Table 7).

**Figure 3:**
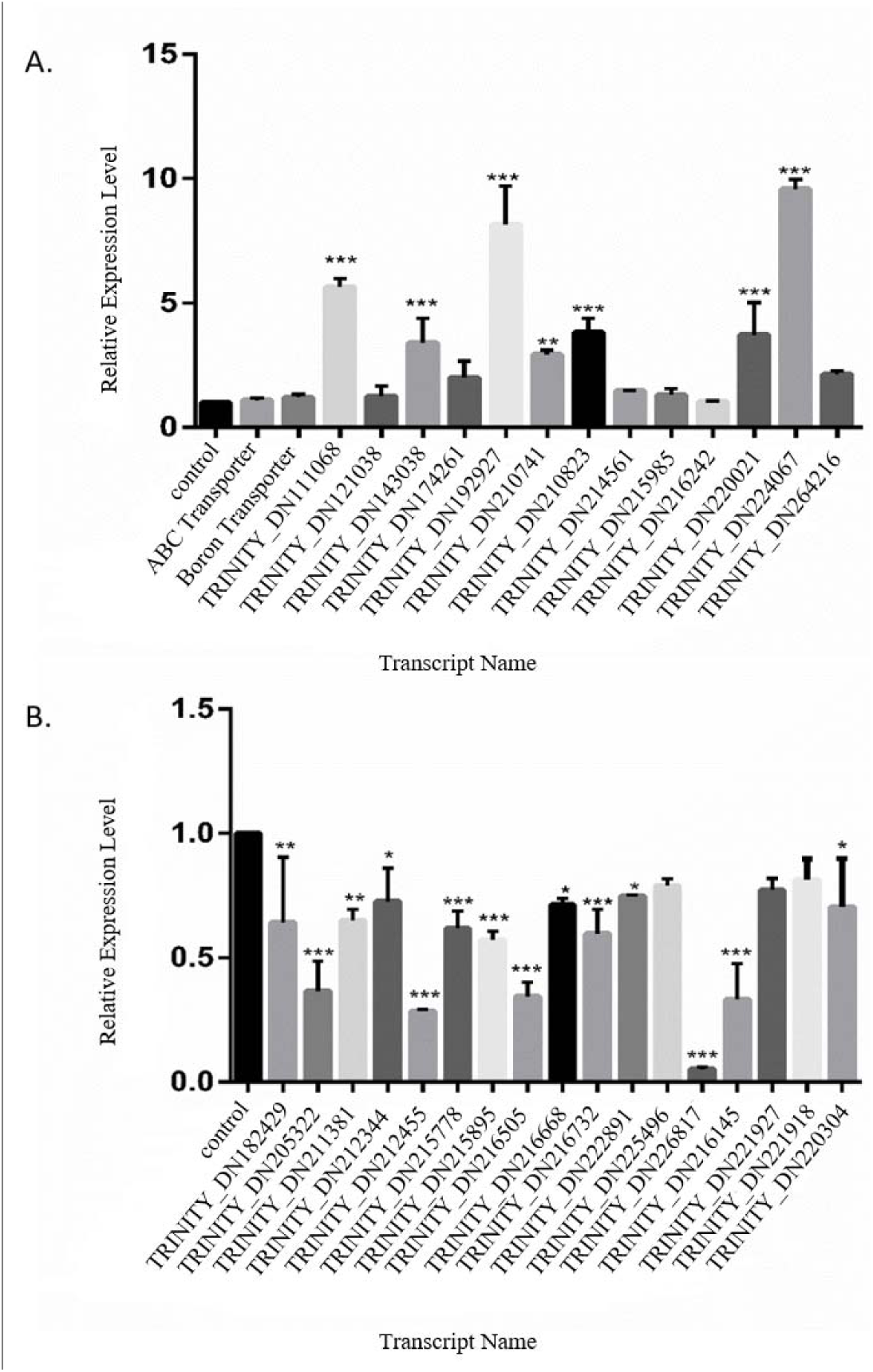
Real-time PCR results of *P. distans* transcriptome for 32 selected transcripts. (A) Fifteen of the transcripts were confirmed to be up-regulated; while 17 were confirmed to be down-regulated (B). Actin was used as internal control and error bars indicate standard deviation of three technical repeats. *** indicates significant difference at p ≤ 0.001, ** indicates significant difference at p ≤ 0.01 and * indicates significant difference at p ≤ 0.05 with respect to control.

### 4. Discussion

Phytoremediation and phytoextraction are clean, simple, cost-effective, environmentally friendly methods for removing toxic elements from soil (Rámilaa et al., 2015). These methods rely on plants such as *P. distans*, an extreme hyperaccumulator of boron. Elucidation of the mechanism(s) by which *P. distans* accumulates and tolerates normally toxic levels of boron may allow development of a faster growing, higher biomass boron hyperaccumulator. This study provides the first broad examination of *P. distans’* transcriptomic response to boron hyperaccumulation. According to the analysis of RNA-Seq data generated in this study, 3312 differentially expressed transcripts were identified and 1652 of them were annotated to 1107 unique proteins with homologs in *A. thaliana* and *O. sativa*. In contrast, 1660 transcripts had no homologs in *A. thaliana or O. sativa*. Thus, these transcripts may be specific to *P. distans* and some may play roles in boron hyperaccumulation, a mechanism which is not present in *A. thaliana* and *O. sativa*.

Plants activate functional and regulatory genes to avoid or tolerate disruptive situations that accompany environmental stresses such as drought, cold, salinity and metalloid accumulation (Shinozaki et al., 2003,Shinozaki and Yamaguchi-Shinozaki, 2007). Many of these genes play roles in basic plant metabolism and are not specific to the stress response. Therefore, this discussion will only focus on stress-related proteins and plant hormones that are differentially regulated under excess boron.

#### Stress-related molecules

Stress-related molecules are known to have roles in the plant’s response to a wide variety of biotic and abiotic conditions (Kobayashi et al., 2014). According to KEGG analysis, certain stress-related molecules were highly active under excess boron as compared to normal boron levels. For example, the *A. thaliana* homolog of UDP-glycosyltransferase (UniProtID: U88A1_ARATH; Transcript Name: TRINITY_DN223147_c5_g2_i1:16-443), which plays a role in flavonoid biosynthesis, was highly up-regulated (Suppl. Table 7). Flavonoids are well-known antioxidants and are important in detoxifying the excess free radicals that plants produce under stress.

Another pathway that was affected by high boron stress was the malate pathway. Activation of this pathway is correlated with abiotic stress (Kumar et al., 2000). Four malate-related *A. thaliana* homolog proteins (UniProtIDs: B3H477_ARATH, FUM1_ARATH, FUM2_ARATH, and MAOP3_ARATH) were down-regulated under excess boron. In contrast, malate dehydrogenase 1 enzyme, which is responsible for converting malate to oxaloacetate, was up-regulated (UniProtID: MDHC1_ARATH) (Suppl. Table 7). This combined increase in malate dehydrogenase 1 and decreased expression of two fumarate hydratases (UniProtIDs: FUM1-2 and B3H477) under boron stress indicates altered regulation in the glyoxylate pathway at malate (Figure 4A) which is in line with previous findings (Beevers et al., 2014). In this pathway, isocitrate is converted to malate via the intermediate molecule, glyoxylate. Glyoxylate and pyruvate can also be formed by a low affinity reaction catalyzed by 4-hydroxy-4-methyl-2-oxoglutarate aldolase (Figure 4B) (Maruyama, 1990). In boron-stressed plants, two 4-hydroxy-4-methyl-2-oxoglutarate aldolases (RRAA2_ARATH and RRAA3_ARATH) were significantly up-regulated (FPKMs: 426.28 and 253.61, respectively) (Suppl. Table 7). Thus, high activity of these aldolases can result in increased glyoxylate which is then converted to malate. Malate can then be transported out of the cell with ions like K+ in order to balance cellular pH, a mechanism that is known to be a response to aluminum tolerance in wheat (Ryan et al., 1995). Thus, these results indicate that malate and related proteins have critical roles in boron hyperaccumulation.

**Figure 4:**
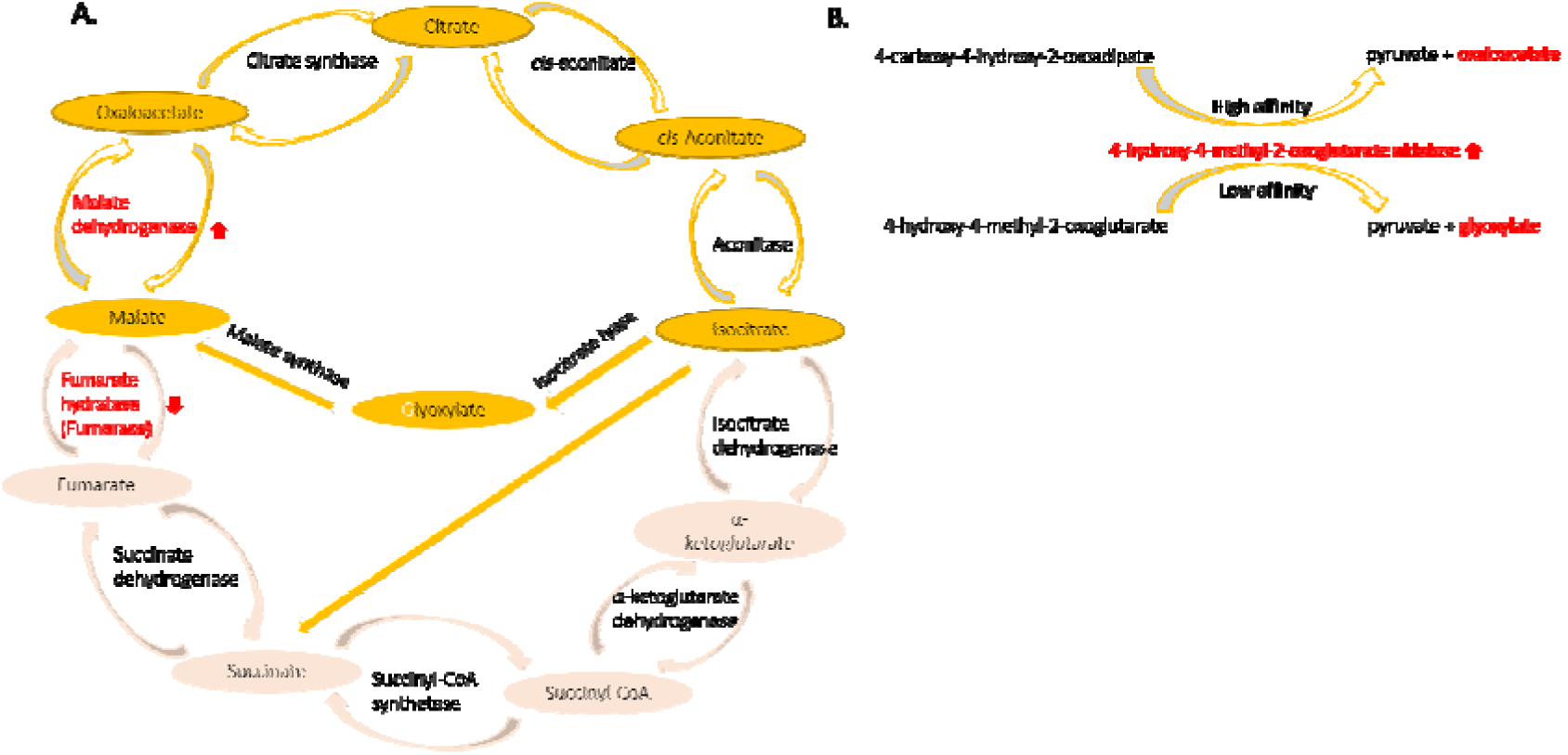
The glyoxalate pathway and B) 4-hydroxy-4-methyl-2-oxoglutarate aldolase reactions. Molecules in red font were affected by boron stress.

Plant cells are enclosed within cell walls which form physical barriers and provide tolerance to turgor pressure (Baskin, 2005). The cell wall consists of polysaccharides, proteins and phenolic compounds (Levy and Staehelin, 1992). Boron is another important structural component of the cell wall as it crosslinks rhamnogalacturonan-II molecules, thereby providing stability. Expansin proteins are also found in the wall and are responsible for cell enlargement by loosening the wall (Cosgrove, 2005). Expansins directly interact with RD22 proteins to mediate cell enlargement (Wang et al., 2012). RD22 proteins contain a BURP domain, an amino acid sequences which is highly conserved in plant species (Xu et al., 2010). As an acronym, BURP comes from: BNM2 (*Brassica napus* microspore-derived embryos protein) (Treacy et al., 1997), USP (*Vicia faba*; unidentified seed protein) (Bassüner et al., 1988), RD22 (*A. thaliana* protein responsive to desiccation) (Yamaguchi-Shinozaki, K. Shinozaki, 1993), and PG1β (*Solanum lycopersicum* β subunit of polygalacturonase isozyme 1) (Zheng et al., 1992). Here we detected, the upregulation of the *A. thaliana* homolog of the RD22 protein (Table 3). In addition to its role in cell enlargement, RD22 was reported to be responsible for increased levels of lignin precursor in cell walls under salt stress (Wang et al., 2012). Under high boron, the excess boron has been shown to interact with lignin precursors in *Pinus radiata* organ cultures (Reid, 2007). Therefore, the up-regulated RD22 homolog in *P. distans* may provide additional lignin precursors, thereby increasing the boron binding capacity of cell walls under stress. Thus, RD22 may be an important component of the hyperaccumulation mechanism in *P. distans*.

#### Transporter proteins

The first identified boron tolerance-related gene was *Bot1* which expresses a putative transmembrane B efflux transporter in barley (Reid, 2007). Many studies on boron efflux transporters showed that the presence of these transporters is controlled by cellular boron concentration. Miwa et al. (2007)observed that under low boron concentrations, AtBOR1 facilitates boron uptake from roots to shoots; while under high boron, AtBOR1 undergoes endocytosis (Miwa et al., 2007; Miwa et al., 2006). These findings indicate that boron concentration affects the dynamics of transporter proteins. A possible explanation is that boron itself interacts with the ribose in mRNA and changes its stability thereby affecting transcription and translation (Pommerrenig, Diehn, & Bienert, 2015). In our study, the *A. thaliana* BOR6 homolog (UniProtID: BOR6_ARATH), a transporter for boron efflux, was up-regulated in *P. distans* (Suppl. Table 7). Since *P. distans* is a hyperaccumulator plant, potential roles of this boron transporter in the cell are: *i)* facilitation of boron transport from root to shoot and *ii)* transport of boron from the cell into the cell wall.

Major intrinsic proteins (MIPs), also known as aquaporin proteins, facilitate the transport of water across biological membranes and are permeable to other small, uncharged molecules such as glycerol, solutes and ions (Quigley et al., 2002; Tyerman et al., 2002; Aharon et al., 2003; Pommerrenig et al., 2015). MIPs are categorized into seven subgroups in the plant kingdom including NOD26-like intrinsic proteins (NIPs) and tonoplast intrinsic proteins (TIPs) transporters (Afzal et al., 2016; Pérez Di Giorgio et al., 2016). NIPs are generally located in the plasma membrane and endoplasmic reticulum; TIPs are localized in the vacuole membrane (tonoplast) and thylakoid inner membrane (Afzal et al., 2016). NIPs are the most divergent plant MIP subfamily with respect to their substrate specificities and amino acid sequences. The NIP aquaporin family is unique to plants and NIPs are selectively permeable to metalloids (Pommerrenig et al., 2015). Specifically, NIP II and NIP III actively transport boron. In our study two types of aquaporins were identified, one of them was a homolog of an *O. sativa* NIP (UniProtID: Q949A7_ORYSA; Transcript Name: TRINITY_DN210823_c0_g3_i1:4-269) and the other was a homolog of an *A. thaliana* TIP (UniProtID: TIP13_ARATH; Transcript Name: TRINITY_DN222891_c3_g1_i1:0-216) (Suppl. Table 7). The *O. sativa* homolog NIP-type aquaporin was up-regulated in *P. distans* under boron stress.Overexpression of an aquaporin could cause tolerance to distinct stress conditions. For example, *PgTIP1* was overexpressed in *A. thaliana* and plants showed increased tolerance to salt and drought stress but decreased tolerance to cold(Peng et al., 2007). Therefore, our results suggest that the *O. sativa* NIP aquaporin is responsible for cellular boron uptake in *P. distans*. On the other hand, the *A. thaliana* TIP homolog was down-regulated indicating that not all aquaporins are involved in boron transport and/or that the vacuole is not an important storage site for boron.

ABC transporters are members of a diverse family. Their susbtrate specificities vary from carbohydrates to ions and some of them are completely specialized to transport ions and solutes. For example, mntABCD is a ABC transporter and selectively permeable for manganese (Baumgart & Frunzke, 2015). In our study, five different *A. thaliana* ABC transporters (ABC-B, C, E, F, I) were differentially expressed under boron stress. All of the ABC transporters were up-regulated with the exception of AB17I_ARATH. The activation of this transporter group may be due to the fact that: *i)* ABC transporters are permeable to a wide variety substrates and some of them may be able to transport boron and *ii)* they can transport boron-binding molecules such as sorbitol. Thus, ABC transporters may be a significant component of the boron hyperaccumulation mechanism in *P. distans*.

Under excess boron, a sulfate transporter was down-regulated in *P. distans* (UniProtID: Q8S317_ORYSA). Sulfate transport mechanisms are divided into four groups. The first is sulfate co-transport with a proton, the second is co-transport with sodium (Na), and the third is sulfate antiport with an anion. The fourth mechanism is ATP-dependent ABC transporter-mediated sulfate uptake (Sze et al., 2012). Sulfate transporters play important roles in response to metal stresses in a metal-specific manner (Kumar et al., 2011). Sulfate transporters are responsible for transport of some oxyanions, such as molybdate and chromate (Appenrothaet al., 2008; Fitzpatrick et al., 2008; Kumar et al., 2011). Kumar et al. (2011)observed that seven sulfate transporters were up-regulated by arsenate and cadmium, ten were up-regulated by chromium and five by lead exposure. However, our study indicated down-regulation of a sulfate transporter in response to boron hyperaccumulation. Because excess boric acid will acidify the cellular environment, less proton pumping is required to maintain cellular pH.

Depending on plant type, sugars (such as sucrose), sugar alcohols (such as mannitol) and amino acids (such as glycine) accumulate under stress conditions and these help plants cope with stress (Taji et al., 2002; Bartels and Sunkar, 2005). Sugar alcohols provide protection from stress by scavenging hydroxyl radicals and/or stabilizing macromolecular structure (Seki et al., 2007). Inositol is a sugar alcohol which also has a role in the biogenesis of the uronosyl and pentosyl units of pectin, hemicelluloses, and related structures in plant cell walls (Loewus and Loewus, 1983; Loewusa and Murthy, 2000). An inositol transporter was up-regulated in *P. distans* (UniProtID: INT2_ARATH; Transcript name: TRINITY_DN172179_c0_g2_i1:0-695,2.5518) under high boron. An increased level of this transporter could be used in the cell to increase the level of inositol available to scavenge reactive oxygen species and/or chelating boron, thereby protecting the cell. Increased cellular inositol may also be required for the synthesis of cell wall pectin and its precursor to bind to the increased amount of boron in the cell wall. The similarity between the inositol and boron transporters of *A. thaliana* (Tanaka and Fujiwara, 2008) suggests a third possibility: the inositol tranporter may also transport boron.

#### Plant hormones

Plant hormones play critical roles in stress conditions by providing physiological and biochemical responses to stress factors (Colebrook et al., 2014). Abscisic acid (ABA) signaling is the central regulator of the abiotic stress resistance pathway in plants (Hubbard et al., 2010). Under drought conditions, plants cope with stress by increasing the amount of ABA causing stomatal closure and reduced transpiration (Cutler et al., 2010). In our dataset, an *A. thaliana* homolog of ABA and related proteins such as phosphatase (UniProtID: P2C16_ARATH; Transcript Name: TRINITY_DN220107_c1_g6_i1:12-396) was down-regulated (Suppl. Table 7). The absence of ABA is associated with increases in cytosolic free calcium (Allen et al., 1999), calcium-dependent proteins (Mustilli et al., 2002) and reactive oxygen species (Murata et al., 2001). According to our findings, calcium-dependent proteins were up-regulated under excess boron, thus, supporting the relationship between ABA and calcium-related proteins. In addition, it is well-known that both calcium and boron are important components of pectic polysaccharides in the cell wall (Matoh and Kobayashi, 1998). Therefore, an increase in boron could trigger up-regulation of calcium levels and calcium-dependent proteins in order to maintain cell wall integrity. Ethylene and ABA are antagonists: ethylene triggers catabolism of ABA (Colebrook et al., 2014). In our analysis, five ethylene response proteins had increased expression under stress (Suppl. Table 7). Thus as expected, the ethylene response was up-regulated while ABA was down-regulated under boron stress.

Another plant hormone that has a role in stress defense is gibberellin (GA). GAs are involved in cell wall loosening in plants, thus, contributing to cell expansion (Cosgrove, 1993; Park et al., 2015). In addition, GAs play roles in plant development and growth, thus they are inhibited under stress conditions to limit growth (Yamaguchi, 2008). For example, GA was reduced under high salinity stress in *A. thaliana* (Magome et al., 2004). Interestingly, in our study, positive regulators of GAs (UniProtIDs: TSN1_ARATH, TSN2_ARATH; Transcript Names: TRINITY_DN151004, TRINITY_DN122991) had increased expression under boron stress (Suppl. Table 7). The excess GAs produced under stress may maintain cell wall looseness as it is known that boron toxicity is usually accompanied by increased cell wall rigidity. Thus, GAs may have a critical role in *P. distans* boron tolerance by preventing cell rigidity, thereby allowing survival and continued growth.

### 5. Conclusion

*P. distans* is a hyperaccumulator, monocot plant and is a non-model organism. Limited knowledge about this species and its hyperaccumulation mechanism led us to use a transcriptomics approach to elucidate stress tolerance and potential hyperaccumulation mechanisms in *P. distans*. Signaling and metabolic pathways in Puccinellia were studied under salinity, alkaline soil and chilling stress conditions (Babao□lu et al., 2004; Padmanabhan et al., 2012; Zhao et al., 2016), therefore, we focused on stress-related molecules, transporters and plant hormones. Strong evidence was obtained that the boron tolerance and hyperaccumulation mechanism of *P. distans* involves alterations in the malate pathway, changes in cell wall components that allow sequestration of excess boron without toxic effects, and at least one putative boron transporter and two putative aquaporins. Transgenic overexpression of this boron transporter and aquaporins in *P. distans* or determination of their subcellular localization will provide deeper information about their direct role in the boron hyperaccumulation mechanism. The next step after confirmation, these genes could be transferred to economically important plants to gain boron tolerance. These results provide new information and insights into the underlying boron stress responsive mechanism in shoots and they are applicaple for agronomy. Additionally, using Puccinellia, itself for phytoremediation and/or phytoextraction will provide new and economical step to remove excess boron from contaminated soil. Therefore this will be useful for soil depollution and for simple boron purification.

